# Drone honey bees (*Apis mellifera*) are disproportionately sensitive to abiotic stressors despite expressing high levels of stress response proteins

**DOI:** 10.1101/2021.08.28.456261

**Authors:** Alison McAfee, Bradley Metz, Joseph P Milone, Leonard J Foster, David R Tarpy

## Abstract

Drone honey bees (haploid males) are the obligate sexual partners of queens, and the availability of healthy, high-quality drones directly affects a queen’s fecundity and productivity of her subsequent colony. Yet, our understanding of how stressors affect drone fecundity and physiology is presently limited. We investigated sex biases in susceptibility to abiotic stressors (cold stress, topical imidacloprid exposure, and topical exposure to a realistic cocktail of pesticides), and we found that drones were more sensitive to cold and imidacloprid exposure but the cocktail was not toxic at the concentrations tested. We corroborated this lack of apparent toxicity with in-hive cocktail exposures via pollen feeding. We then used quantitative proteomics to investigate protein expression profiles in the hemolymph of topically exposed workers and drones, and we show that drones express surprisingly high levels of putative stress response proteins relative to workers. Drones apparently invest in strong constitutive expression of damage-mitigating proteins for a wide range of stressors, yet they are still sensitive to stress when challenged. The robust expression of stress-response proteins suggests that drone stress tolerance systems are fundamentally rewired relative to workers, and their susceptibility to stress depends on more than simply gene dose or deleterious recessive alleles.

## Introduction

High quality male honey bees (*Apis mellifera* drones) are essential for supporting adequate mating of queens, whose longevity depends on the number and quality of sperm acquired during nuptial flights. Despite being critical players in honey bee reproduction, factors affecting drone quality are understudied topics (reviewed recently by Rangel *et al*.^1^). Limited existing research has shown that some pesticides^2-5^ and extreme temperatures^6-9^ negatively impact drone fecundity, but generally, little is known about drone abiotic stress tolerance and their stress-mitigating responses.

Drones have significantly lower tolerance thresholds to heat stress relative to workers^6^. This phenomenon may be in part explained by the haploid susceptibility hypothesis, which states that haploid individuals are more susceptible to stressful conditions, such as pathogenic infections and abiotic stressors, since they have no opportunity for heterozygous buffering of deleterious recessive alleles^10^. Baer *et al*. found that 77% of drones died after exposure to 42 °C for 4 h^7^, and McAfee *et al*. found that 50% of drones died after exposure to 42 °C for 6 h, whereas only 2% of workers perished under the same conditions^6^.

However, the haploid susceptibility hypothesis is not consistently supported when it comes to pathogenic infections^11^. While investigations on honey bee male susceptibility to *Nosema*^12^, as well as immunocompetence of leafcutter ants (*Atta colombica*)^13^, wood ants (*Formica exsecta*)^14^, and buff-tailed bumble bees (*Bombus terrestris*)^15^ support the haploid susceptibility hypothesis, research on *B. terrestris* male susceptibility to *Crithidia* does not^11^. Furthermore, sex biases in pathogen susceptibility exist for both male and female insects, depending on the species, even those without haplo-diploid sex determination systems^16-18^. This suggests that immunocompetence is a highly complex trait with many interacting factors^12^, such as underlying infections and even abiotic stressors^19^, obscuring the true patterns of susceptibility.

Neonicotinoid pesticides have been described as “inadvertent insect contraceptives,” owing to evidence that thiamethoxam and clothianidin can reduce drone fecundity and lifespan during colony-level exposures when colonies were fed pollen patties containing low concentrations of the insecticides (< 5.0 ppb)^3^. Furthermore, topical exposure to 2 µl of 20 ppb imidacloprid, another neonicotinoid, has been shown to reduce viability of sperm stored within queens^20^. Friedli *et al*. found that thiamethoxam and clothianidin had a greater impact on developmental stability of drones compared to workers, and attribute this pattern to be driven in part by male haploidy^2^.

Experiments documenting effects of exposure to specific classes of pesticides are important; however, since drones do not forage, they are most likely to encounter more complex pesticide mixtures that accumulate in various hive matrixes (*e*.*g*., wax, pollen, honey). In a survey of commercial honey bee colonies in the U.S., Traynor *et al*. documented residue data for pesticides, herbicides, and fungicides present in beebread, wax, and other hive components^21,22^. These data offer a realistic reference point for investigating effects of compound cocktails in realistic abundances and relative proportions. While high residue concentrations within hive matrices have been linked to queen failure^21,23^, which was likely driven by indirect effects on worker jelly secretions rather than direct effects on the queen^24,25^, the impact of hive residues on drone physiology has not yet been investigated.

Here, we aim to investigate drone and worker tolerances to abiotic stressors, focussing mainly on pesticide exposure. We confirm drone susceptibility to imidacloprid, and we further investigate the impacts of drone exposure to pesticide cocktails (based on data in Traynor *et al*.^21^) through topical applications as well as supplemental hive treatments. Finally, we investigate drone and worker stress responses to topical pesticide applications (control, imidacloprid, and cocktail treatments) through proteomics analysis of the hemolymph. Our data suggest that drones have surprisingly strong baseline expression of putative stress response proteins, contrary to our expectations, causing us to re-evaluate exactly why drones, but not workers, are so intolerant to abiotic stress.

## Results

### Sex biases in survival across abiotic stressors

We and others have previously reported sex biases in heat tolerance, with upwards of 50% of drones perishing after exposure to 42 °C for 6 h, whereas only 2% of workers died after the same treatment^6,7^. To determine if this sex bias exists across other abiotic stressors, we also compared worker and drone sensitivity to cold (4 °C, acting as a positive control for mortality), imidacloprid (1, 10, and 100 ppm, topical exposure), and a cocktail of the nine compounds frequently found in wax (as described in McAfee *et al*., recipe derived from Traynor *et al*.^21^; administered at 0.33x, 2x, and 10x, where x is the median concentration in wax). Baseline drone survival in the negative control groups ranged from 91 to 93% in all three experiments, whereas worker survival was 100%. As expected, we found a strong sex bias in cold tolerance; no workers perished in the experiment, but drone survival counts were significantly affected by cold exposure (z = -3.6, df = 45, p = 0.00031), with 76% and 92% of drones perishing after 2 and 4 h exposures at 4 °C, respectively (**Figure 1a**). Worker survival was also not affected by imidacloprid exposure at the tested doses and replication (z = -0.01, df = 115, p = 0.99). Drone survival, however, was significantly affected by increasing imidacloprid dose (z = -1.99, df = 115, p = 0.047; **Figure 1b**). These are highly unrealistic exposure scenarios and are strictly employed to investigate sex-biases. No appreciable drone or worker mortality was observed with exposure to any cocktail dose (workers: z = -0.003, df = 76, p = 1.0; drones: z = -0.079, df = 81, p = 0.94), indicating that the agrichemical matrix commonly found in wax has low contact toxicity to both male and female bees (**Figure 1c**).

**Figure 1.**
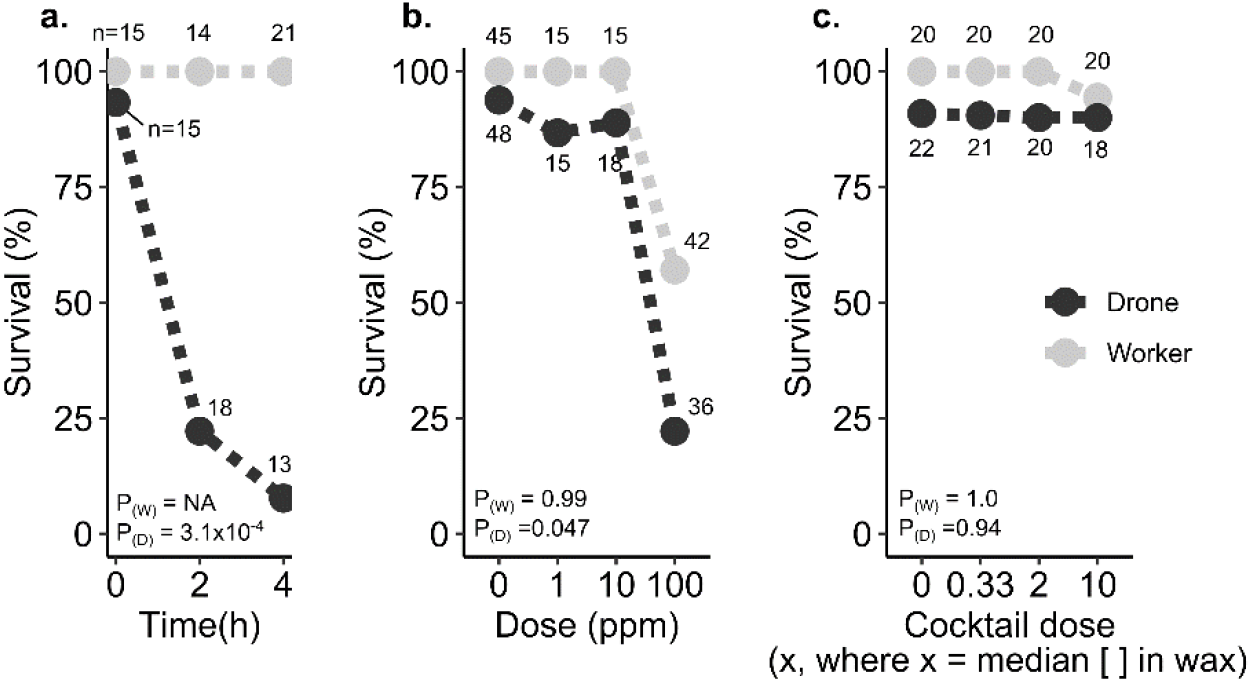
Sex biases in survival during temperature and pesticide stress challenges. Drones and workers (all five days old at the beginning of the experiment) from three different colony sources were marked and kept in JZ-BZ queen cages with candy (one subject with five other companion workers of unknown ages in each cage). Statistical differences were evaluated using a ⍰^2^ test on survival count data. a) Baseline survival rates of the negative control group (2 µl topical acetone treatment for ‘cocktail’ and ‘imidacloprid’ groups, and room temperature incubation for the ‘cold’ group) in the three different experiments after two days. b) Survival (normalized to baseline) of drones and workers two days after exposure to different durations of cold stress (4 °C). c) Normalized survival of drones and workers topically exposed to different concentrations of imidacloprid. d) Normalized survival of drones and workers topically exposed to different concentrations of a pesticide cocktail of compounds commonly found in wax (see McAfee *et al*.^26^ for the recipe, and Traynor *et al*.^21^ for the supporting data).

### Equivocal effects of in-hive cocktail exposures on drone survival, body size, and fecundity

Since the agrichemical cocktail is the exposure that drones are most likely to experience in a managed setting, and prior evidence suggests that exposure through pollen poses a greater hazard than wax^25^, we aimed to corroborate our negative results of topical cocktail exposure with hive exposures via pollen patties, targeting either drone adults (experiment 1) or drone larvae (experiment 2). In the first experiment, we found that adult drones banked in colonies fed control pollen patties actually had greater mortality than colonies fed patties containing the pesticide cocktail (⍰^2^_1_ = 19.8; p < 0.0001). However, this trend was only significant for drones from one of the two source colonies (source one: ⍰^2^_1_ = 0.0846; p = 0.771; source two: ⍰^2^_1_ = 38.2; p < 0.0001) and this is further confounded by significant differences among mortality within the banks within each treatment/source group (minimum ⍰^2^_2_ = 11.7, p = 0.003; **Figure 2a**).

**Figure 2.**
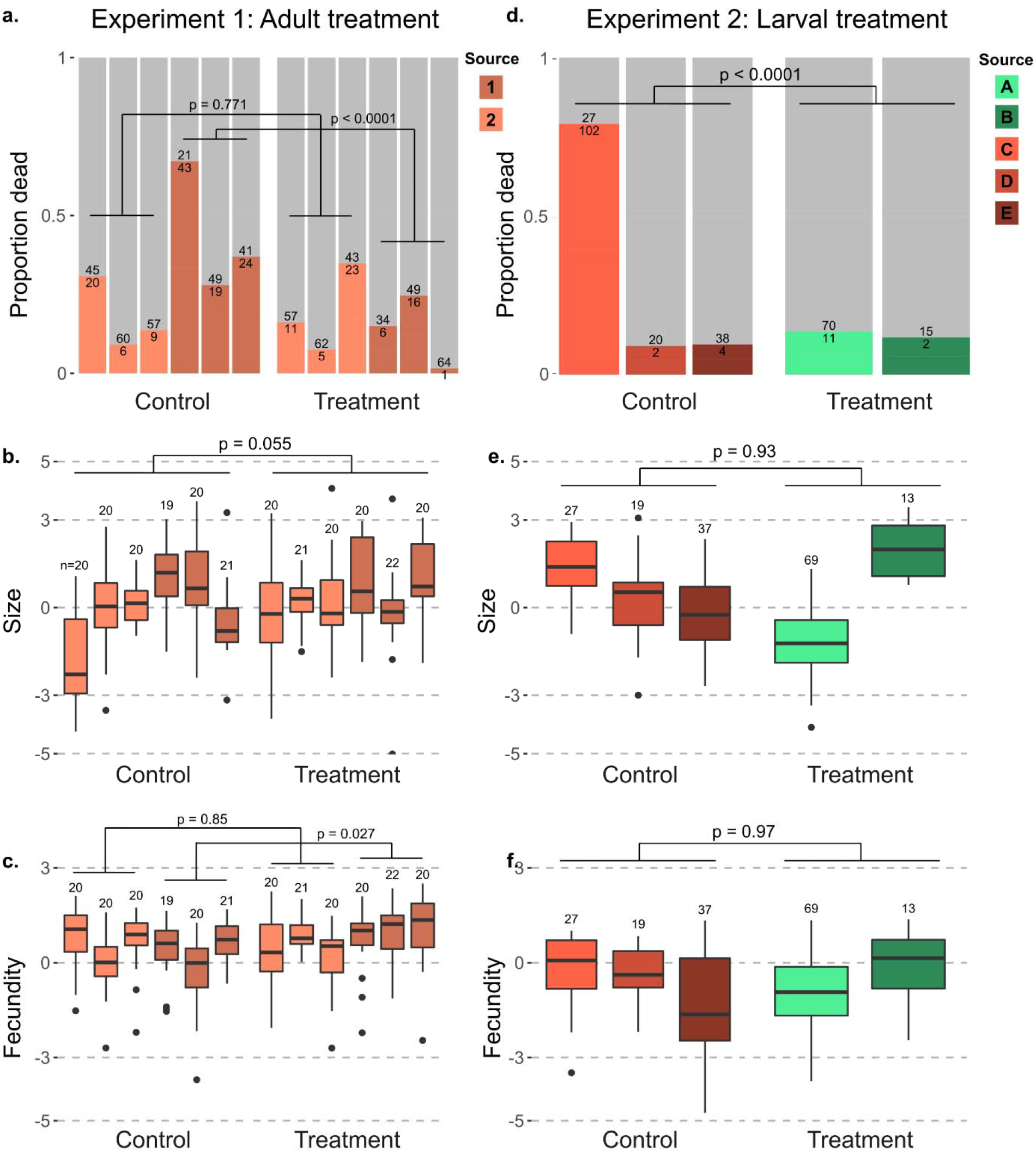
No consistent effects of hive-level cocktail treatments via pollen on drone development or adult fostering. Drones were exposed to colony-level pollen-delivered pesticide cocktail either as adults (experiment 1) or as larvae (experiment 2). Unique drone source colonies are differentiated by color. Number of individuals included in each group is displayed over their respective bar or boxplot. Mortality data are reported as proportion dead in color, with live proportion in grey. Mortality differences were evaluated with ⍰^2^ tests and size and fecundity were evaluated using linear mixed models (see methods for specific models). (a-c) Mortality, size, and fecundity of drones from different source colonies reared in untreated colonies but fostered in either treated and untreated colonies as adults. (d-f) Mortality, size, and fecundity of drones from different source colonies which were reared through development by treated and untreated colonies, but fostered in untreated colonies as adults.

We next investigated effects of colony source, bank treatment, and their interaction on drone fecundity (see Methods for a definition of this parameter). We found that drone size differed only across colony source (linear mixed model, t_237.1_ = 3.72; p = 0.0002), with neither an effect of treatment (t_14.4_ = 2.09; p = 0.055) nor source by treatment interaction (t_237_ = -1.56; p = 0.12) or bank colony (likelihood ratio = 1.23; p = 0.267; **Figure 2b**). Testing the same model for drone fecundity revealed a significant treatment by source interaction (t_237_ = 2.99; p = 0.003), so we then tested each source separately. Fecundity of drones from source one did not differ because of treatment (t_6_ = -0.197; p = 0.85) or adult bank (likelihood ratio = 3.63; p = 0.057). However, fecundity of drones from source two differed due to treatment (t_6_ = 2.90; p = 0.027) but not adult bank (likelihood ratio = 1.18; p = 0.28), with the trend of drones banked in treated colonies actually having higher fecundity (**Figure 2c**). That this is not visibly obvious is indicative of the equivocal nature of these findings.

We expected that worker care at the larval stage could have a greater or more consistent effect on drones, as has been observed for queens previously^24,27^. We therefore also investigated effects of colony cocktail exposure via pollen patties on the quality of drones they reared. We found that drone emergence numbers were not significantly different across treatment (F_1,25_ = 3.08; p = 0.09) though colonies ranged widely in drones produced (0-120 individuals). Drone mortality followed a similar trend as in experiment 1 with a strong trend toward higher mortality among the control treatment (⍰^2^_1_ = 47.0; p < 0.0001) driven by strong differences in mortality among control sources (⍰^2^_2_ = 84.3; p < 0.0001) rather than treatment sources (⍰^2^_1_ = 1.3×10 ^-30^; p = 1) and a clear confounded factor of near total mortality of a single control source (**Figure 2d**). Drones differed in size based solely on the colony source (likelihood ratio = 68.9; p < 0.0001) rather than larval treatment (t_5_ = -0.09; p = 0.93; **Figure 2e**). The results were similar for drone fecundity, with significant differences due to source (likelihood ratio = 9.2; p = 0.002), but not larval treatment (t_5_ = 0.04; p = 0.97; **Figure 2f**).

### Overlap between sex-biased protein expression and proteins expressed by drones in response to pesticide treatment

Despite finding no clear effect of the pesticide cocktail treatments to whole colonies, sex-biased tolerance was apparent for the other stressors tested with topical applications. To investigate the molecular origin of these differences, we performed differential protein expression analysis on hemolymph from workers and drones exposed to the control (acetone), cocktail (10x), or imidacloprid (10 ppm) treatments. This is not meant to investigate effects of realistic exposures (a 10 ppm topical imidacloprid exposure is unrealistically high); rather, the purpose is to determine what proteins are involved in sex-biased stress tolerance (we did not examine cold-stressed drones because too few drones survived the treatment to analyze). We identified 1,452 protein groups in total (1% FDR), but after filtering out proteins without at least three identifications in each experimental group, 654 proteins groups were quantified. Of those, 188 were differentially expressed between drones and workers, and 34 were differentially expressed between control drones and imidacloprid-treated drones (**Figure 3a and b**, all at 5% FDR, Benjamini-Hochberg correction). No differences were identified in any of the other pairwise comparisons, including cocktail treatments relative to controls, further supporting that the cocktail is not hazardous to drones at these doses. Of the 34 proteins differentially expressed between imidacloprid- and acetone-treated drones, 18 were also differentially expressed between drones and workers (**Figure 3c**-**e**).

**Figure 3.**
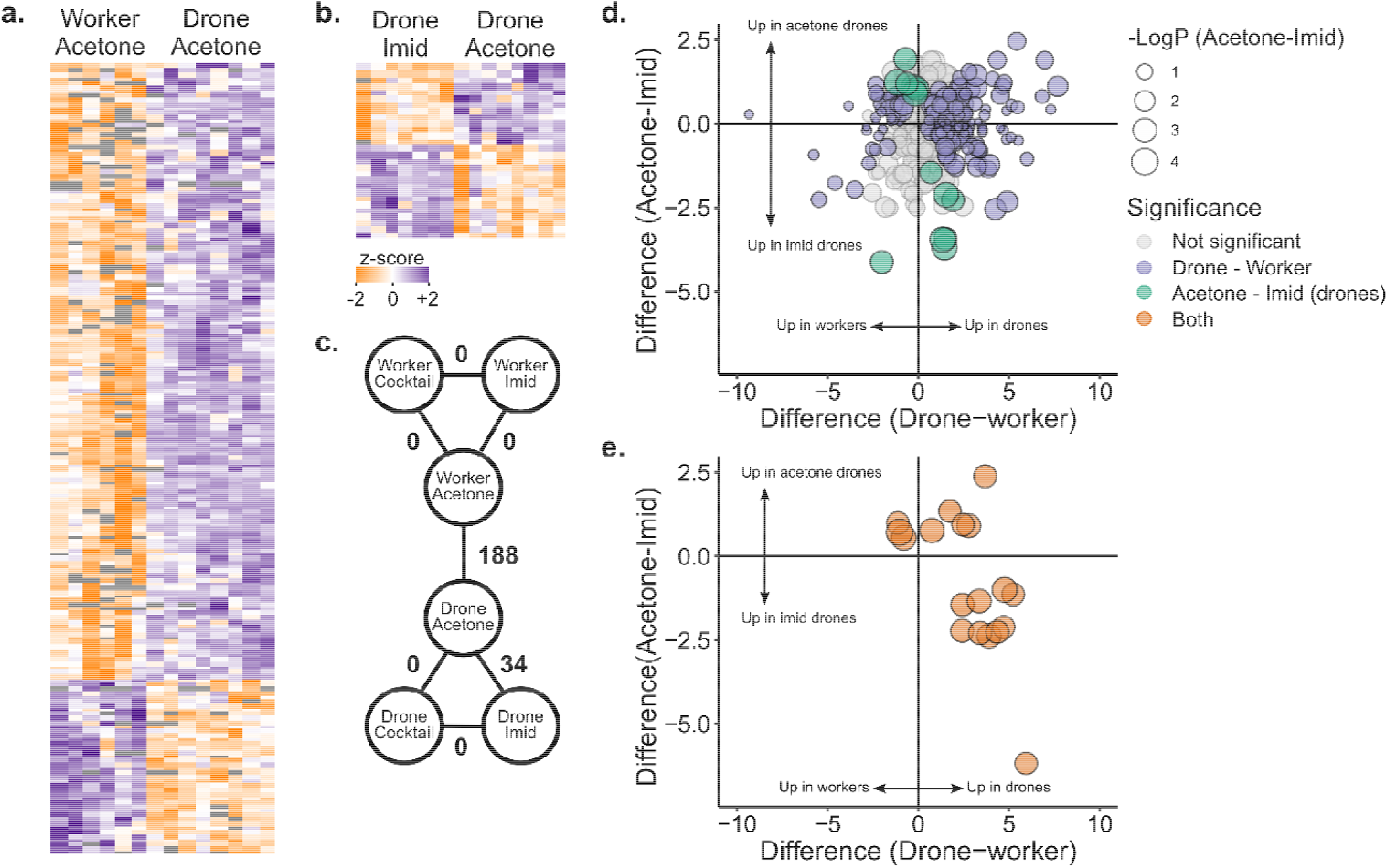
Quantitative proteomics on surviving drone and worker hemolymph after topical exposure to pesticides. Workers and drones exposed to the pesticide cocktail (2 µl of 10x solution in acetone, where x = the median concentration in wax), imidacloprid (2 µl of 1 ppm solution in acetone), and a negative control (2 µl of acetone). a) Label-free quantitative proteomics results comparing negative control drones and workers. Each row is a protein and each column is a sample. Only significant differences (5% FDR, Benjamini Hochberg correction) are depicted. b) Proteins differentially expressed comparing imidacloprid-exposed drones to controls. c) Summary of statistical analyses comparing different groups. Numbers indicate the number of proteins differentially expressed at 5% FDR. d) Summary of significance and direction of change in expression for proteins differentially expressed in drones vs. workers (purple), and imidacloprid vs. control drones (green). Circle size is proportional to −log(p value) for the drone imidacloprid vs. acetone comparison. e) Proteins differentially expressed in both statistical comparisons. Circle size is proportional to −log(p value) for the drone imidacloprid vs. acetone comparison.

We hypothesized that drones might be disproportionately sensitive to abiotic stressors if they are unable to launch an adequate stress response (e.g., detoxification enzymes). We therefore expected that putative stress response proteins would be expressed at lower levels in drones relative to workers. Interestingly, we observed the opposite trend. All but two of 17 putative stress response proteins, including proteins linked to detoxification, oxidative stress, immunity, and heat-shock proteins, were upregulated in drones relative to workers (**Figure 4a**; 5% FDR, Benjamini-Hochberg correction). Among these were three heat-shock proteins (HSP cognate 3, HSP beta 1, and 97 kDa HSP, corresponding to NP_001153524.1, XP_003251576.1, and XP_006561225.1, respectively), which were all expressed more highly in drones than workers, but were not differentially regulated by pesticide exposure (**Figure 4b**).

**Figure 4.**
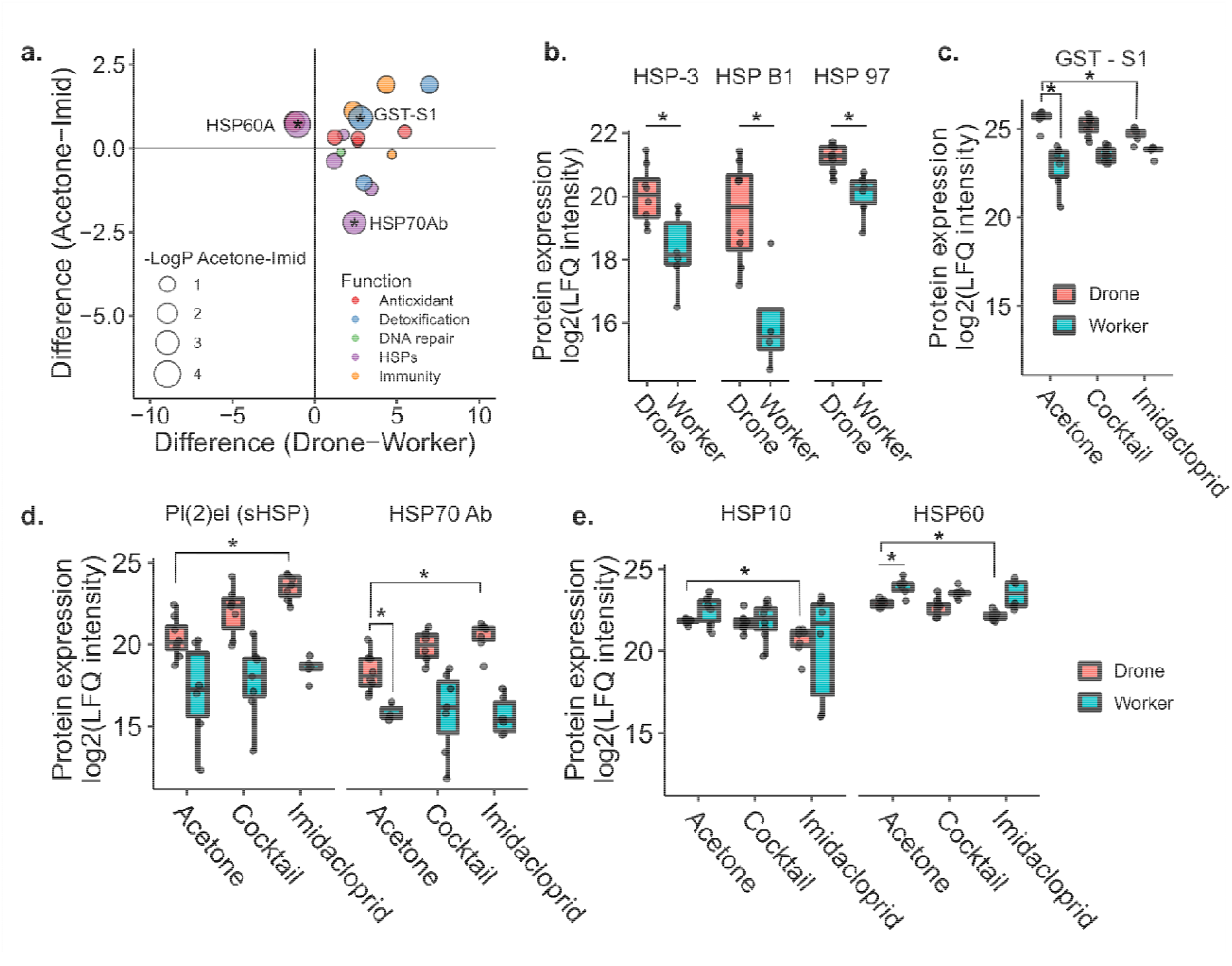
Differentially expressed proteins linked to stress responses. a) Proteins differentially expressed between drones and workers which have functions linked to oxidative stress, detoxification, DNA repair, the heat-shock response, and immunity. Asterisks indicate that the protein was also differentially expressed in drone imidacloprid vs. control comparisons. b) Patterns of expression for HSP cognate 3, HSP beta 1, and 97 kDa HSP in drones and workers. Asterisks indicate that the comparison was significant at 5% global FDR (Benjamini Hochberg). c) Pattern of expression of an uncharacterized protein best matching glutathione-S-transferase S1 in *Drosophila melanogaster* across pesticide treatments. Asterisks indicate that the comparison was significant at 5% global FDR (Benjamini Hochberg). d) Patterns of expression of protein lethal (2) essential for life (Pl(2)el) and HSP70 Ab across pesticide treatments. Asterisks indicate that the comparison was significant at 5% global FDR (Benjamini-Hochberg). e) Patterns of expression of 10 kDa HSP and HSP60, mitochondrial HSPs which are known to physically interact in a 1:1 stochiometric ratio. Asterisks indicate that the comparison was significant at 5% global FDR (Benjamini Hochberg).

An uncharacterized protein (XP_026295805.1) with high sequence homology to *Drosophila melanogaster* glutathione-S-transferase S1, a detoxification enzyme, was also upregulated in drones relative to workers, and downregulated in imidacloprid-treated drones relative to the negative control (**Figure 4c**). Furthermore, glutathione-S-transferase S4 (XP_006560566.1) was one of the top three upregulated proteins in drones relative to workers, with a log2(fold change) of 6.97. HSP70 Ab (NP_001153544.1) was differentially expressed in both sex and pesticide exposure comparisons, but, like pl(2)el (XP_006568238.2; another HSP), expression increased with imidacloprid treatment for drones, but not workers (**Figure 4d**). HSP60 (XP_392899.2) was one of the two putative stress response proteins that was downregulated in drones relative to workers, and it and its binding partner (HSP10; XP_624910.1) were both further downregulated with imidacloprid treatment (**Figure 4e**).

Drones appear to exhibit a robust suite of stress response proteins even in the absence of temperature or pesticide stress, and we hypothesized that a dramatic mobilization of resources would be required to sustain these basal expression levels. Hexamerins are well-known amino acid storage proteins that are highly abundant in larval hemolymph and are thought to be catabolized to support metamorphosis, when the developing bee is unable to feed^28,29^. We expected that adult drones would have low levels of hexamerins in their hemolymph, in order to support the energetically costly maintenance of high basal levels of proteins involved in stress responses. There are four major honey bee hexamerins: hex70a, hex70b, hex70c, and hex110^29^. Only hex70a (NP_001104234.1) and hex110 (NP_001094493.1) were quantified in our dataset. Whereas hex70a did not exhibit sex biased expression patterns, hex110 was actually the most strongly differentially expressed protein between drones and workers, with significantly lower abundance in drones (p = 0.000244, q = 0.00296, t = -5.32; **Figure 5a**), which supports our hypothesis. After hex110, the top 4 proteins exhibiting the strongest sex biased expression were serpin88Ea (XP_026298978.1), trypsin 1-like (XP_026301257.1), inositol-3-phosphate synthase (XP_623377.1), and adenylate kinase (NP_001164443.1; **Figure 5b**).

**Figure 5.**
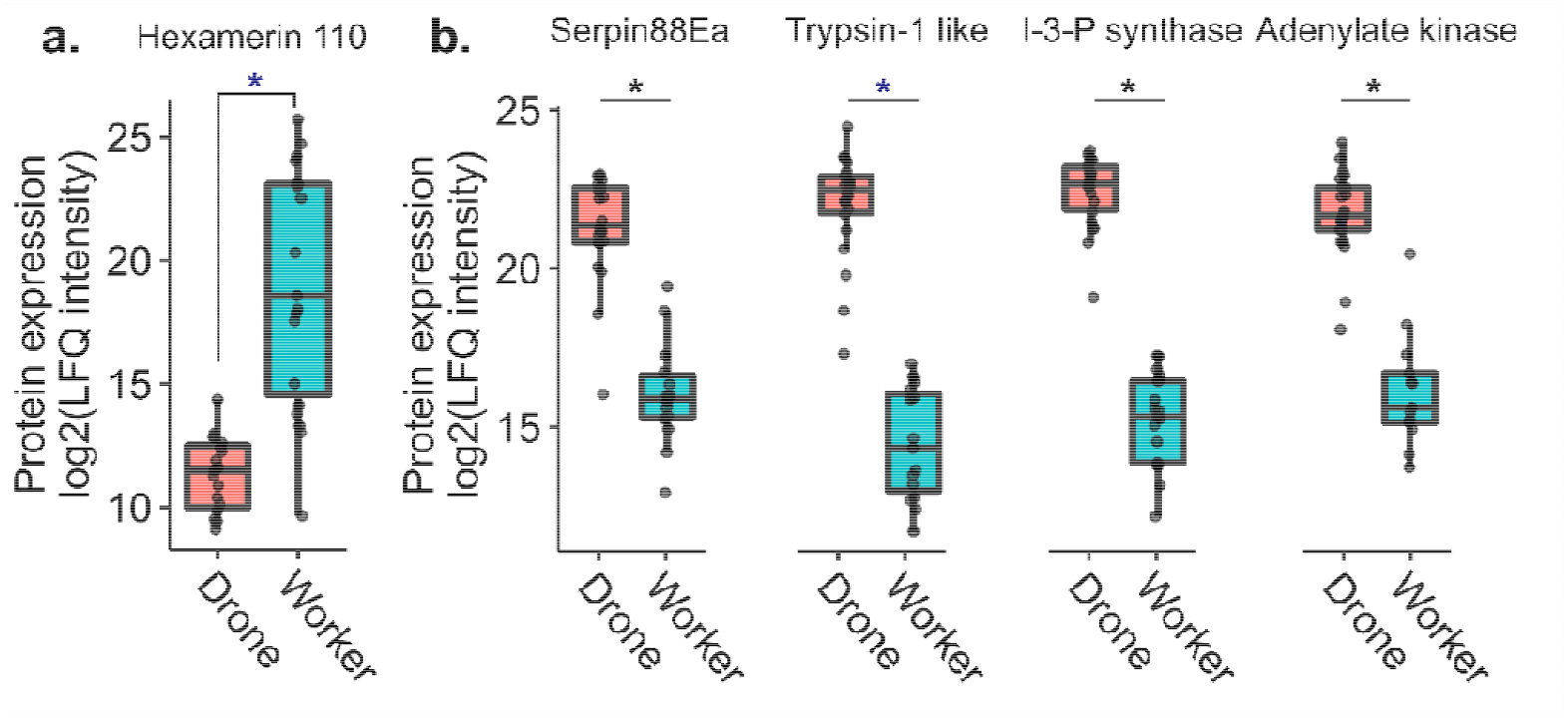
Expression of the top five most strongly differentially expressed proteins, in terms of fold-change, between drones and workers. Expression of these proteins were not linked to pesticide exposure so the exposure data are included in the drone and worker categories. Asterisks indicate that the comparison was significant at 5% global FDR (Benjamini Hochberg). a) Hex110, a conserved amino acid storage protein, was the most strongly differentially expressed protein overall, with very low levels present in the drones. b) The top four other proteins were all more abundant in the drones, including a serine protease inhibitor (serpin88Ea), a serine protease (trypsin-1 like), inositol-3-phosphate synthase, and adenylate kinase.

Because our experimental design did not include untreated controls, it is possible that drones appear to have high baseline levels of stress response proteins relative to workers simply because they have a stronger response to acetone (the negative control). To determine if this could be the case, we compared hemolymph proteomes from age-matched, untreated drones and workers (n = 14 each) sampled from three different hives. Of the 483 proteins that were differentially expressed (out of 988 proteins quantified after filtering), we found that the data largely agree with the findings above. Namely, glutathione-S-transferases were again significantly upregulated in drones (**Figure 6a**), hex110 was again strongly downregulated in drones whereas inositol-3-phosphate synthase was upregulated (**Figure 6b**), and HSPs were largely also upregulated in drones with the exception of HSP cognate 3 and one of the three proteins named pl(2)el (**Figure 6c and d**). These results are consistent with our initial findings and suggest that the results are not an artefact of acetone treatment.

**Figure 6.**
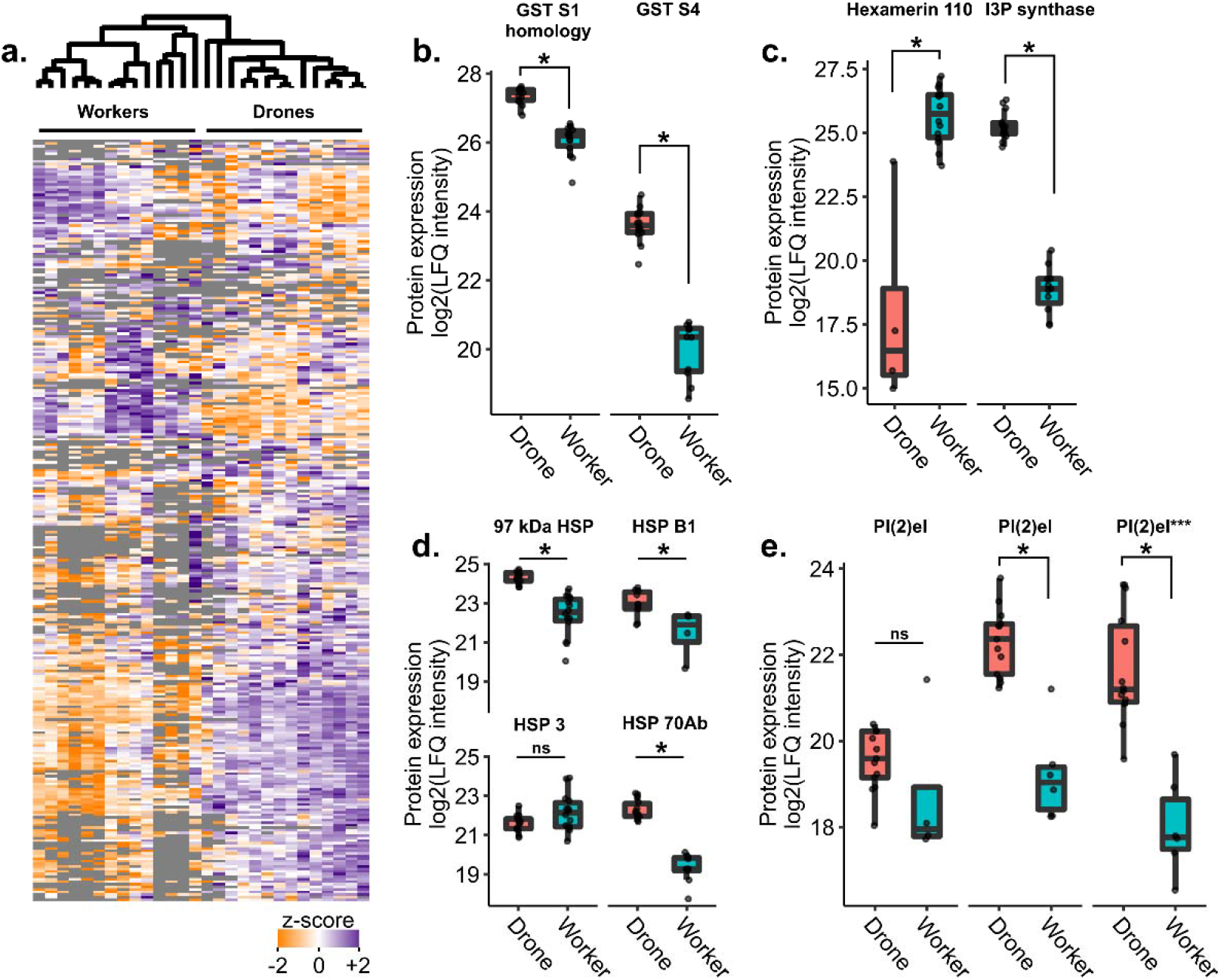
Comparing untreated drones and workers. a) Quantitative proteomics identified 2,089 proteins in the hemolymph of n = 14 drones and n = 14 workers, with 988 quantified after quality filtering. b-e) 483 proteins were differentially expressed (two group comparison, Benjamini-Hochberg correction at 5% FDR), with proteins of interest highlighted here. e) Several proteins share the name protein lethal(2) essential for life-like (pl(2)el), and the specific pl(2)el, which was depicted in Figure 4d is highlighted by three asterisks here.

## Discussion

The haploid susceptibility hypothesis broadly states that haploid individuals are more susceptible to stressors because their haploid state makes it more likely for them to possess deleterious alleles with no opportunity for compensation^2,10^. While we did not examine the presence of mutations in these drones, we did show that adult haploid honey bees (drones) are indeed more susceptible to cold stress and pesticide stress than diploid worker bees, similar to heat stress, as has been demonstrated previously^6,7^. We also further investigated the impact of drone exposure to a realistic agrochemical cocktail via pollen to whole colonies and found no consistent effects of pesticide treatment on adult drones or larvae, indicating that this mixture either does not have appreciable toxicity or that drones are buffered from exposure by indirect feeding through workers. Finally, we identified a surprising general trend for drones to express higher levels of putative stress response proteins compared to workers — findings consistent with the trends observed in the existing honey bee protein atlas comparing sexes, castes, and tissues^30,31^ — and some of the same proteins were also differentially regulated in response to pesticide stress.

This strong expression of stress response proteins was in contrast to hex110 expression, which is a major amino acid storage protein^29^ that was massively downregulated in drones relative to workers. We tentatively propose that mobilization of hex110 may provide the resources needed for drones to express significantly higher constitutive levels of stress response machinery compared to workers, and that this scenario may represent a sacrifice (depletion of hex110 stores) for the sake of a short-term gain (high constitutive expression of stress response proteins). This hypothesis will require further testing, such as with hex110 knock-down experiments, in order to concretely ascertain.

Our results suggest that the haploid susceptibility hypothesis does not fully explain the general sensitivity of drones to stress, since, from a protein abundance standpoint, the drones’ repertoire of proteins that help mitigate damage due to stressors is surprisingly robust. Nor does gene dose, *per se*, explain the trends observed, since drones have half the gene dose as workers yet express higher basal levels of stress response proteins. Rather, we suggest that the drone’s stress response may be a result of sex-specific rewiring of the stress responses, such that most of their amino acid reserves are constantly mobilized to support high baseline levels of proteins that help mitigate damage due to short-term stress. Indeed, drones have concentrations of free amino acids in the hemolymph that are over three times higher than in workers of the same age^32^. This suggests that, rather than being stored as hexamerins, these resources are mobilized, perhaps to support other sex-specific protein expression.

It is somewhat confusing, then, for drones to so easily die in the face of challenging conditions, if their stress response is already primed. One explanation is that, while they generally express high levels of stress response proteins, they have no further amino acid or energy reserves to amplify that response, should a severe intensity or duration of stress be encountered. We suggest that the drone’s investment in high baseline expression proteins linked to oxidative damage, detoxification, temperature stress, DNA damage, and immunity enables them to combat a wide range of mild stressors, but leaves them unable to launch a stronger response to deal with intense, specific stressors. Another explanation is that there may be underlying qualitative differences in drone stress proteins relative to workers. For example, despite finding an increased abundance of the glutathione-S-transferase in drones when compared to workers, drone glutathione-S-transferases may have a reduced detoxification activity towards pesticides. Qualitative differences in honey bee detoxification proteins have been previously reported and it has been found that enzyme abundance does not necessarily correlate with detoxification activity in honey bee workers^33^. Similarly, large qualitative differences in another putative detoxification enzyme, esterase, have been identified in worker larvae from different breeding stocks while simultaneously finding no differences in esterase abundances^27^.

Conversely, these results also raise the question of how workers are so stress tolerant, in terms of survival, without launching an equally robust stress response. Indeed, we identified no differentially expressed proteins comparing workers treated with imidacloprid to controls. Since we only quantified 654 protein groups, out of 1,452 identified proteins and still more proteins which exist below our limit of detection, it is possible that important stress response proteins were simply not quantified in our dataset. However, we still expected to see at least some sign of a stress response. An alternate explanation is that, since these bees were euthanized 2 full days after experiencing the stress, it is possible that workers are more efficient in their stress response than drones, and have already both launched and reversed their stress response. This may be done through differential expression, or in a manner mediated by post-translational modification of proteins to modulate the proteins’ specific activities. These explanations would be consistent with the ability for workers to rapidly and efficiently mobilize, then shut down, a specific stress response as conditions change.

While drones expressed higher levels of many heat-shock proteins (HSP beta 1, HSP cognate 3, 97 kDa HSP, protein lethal(2) essential for life, and HSP70 Ab), as well as a putative Glutathione-S-transferase, they expressed lower levels of HSP60 compared to workers. They also tended to express lower levels of HSP10 (the binding partner of HSP60)^34^. HSP60 and HSP10 assist with folding proteins imported to the mitochondria and prevent protein aggregation^34,35^. HSP10 and HSP60 are further downregulated with imidacloprid treatment, indicating that imidacloprid exposure may make drones even more susceptible to temperature stress than they already are. Determining the additive and synergistic effects of multi-stressor drone exposure as well as indirect effects of drone exposure on queens (such as work done by Kairo *et al*.^36,37^ and Bruckner *et al*. (personal communication)) is an important study area to broaden.

Among the proteins most strongly differentially regulated between drones and workers were serpin 88Ea, trypsin-1 like, inositol-3-phosphate (I-3-P) synthase, and adenylate kinase (**Figure 5**). This is puzzling, as serpin88Ea is a serine protease inhibitor and trypsin-1 like is a serine protease; therefore, their strong co-expression appears to be an inefficient use of resources. However, whether trypsin-1 like is actually a target of serpin88Ea is unknown. In *Drosophila*, serpin88Ea is a negative regulator of the toll immune response via inhibition of spaetzle processing enzyme^38^, and in honey bees, serpin88Ea has been linked to stored sperm viability^39^; therefore, it is possible that these two proteins are actually involved in different processes. I-3-P synthase, however, is an enzyme known to become upregulated with abiotic stress in the Eastern honey bee, and which in turn regulates antioxidant enzymes such as superoxide dismutase and glutathione-S-transferase^40^. Ni *et al*. ^*40*^ found that knockdown of *A. cerana* I-3-P synthase subsequently inhibited antioxidant enzyme expression, for example. Given that our data shows that drones consistently upregulate other stress response proteins, is likely that the enzyme has a similar function in Western honey bees, too. Finally, adenylate kinase plays an important role in regulating cellular energy homeostasis^41^. Its strong upregulation in drones may further point to drones operating on the margins of energy expenditure.

The in-hive drone exposures to a pesticide cocktail were meant to further investigate the null result we obtained when topically exposing drones to different concentrations of a pesticide blend. We reasoned that, similar to previous work conducted on queens and royal jelly production^24,25^, an oral pollen exposure may affect drones where a direct topical exposure does not, potentially via altered worker care or jelly secretions. However, we did not observe consistent effects of pesticide treatment on drone size or fecundity, whether the colonies were exposed during the drones’ development or adulthood. Rather, we observed that the drone source colony had the most pronounced effect on these quality parameters, indicating that, at the very least, any colony level effects of cocktail exposure that might exist are far outweighed by other natural colony parameters.

## Conclusion

We found that drones are more sensitive, in terms of survival, to cold stress and imidacloprid exposure compared to workers, and that drones exhibit surprisingly strong constitutive expression of putative stress response proteins. These proteins have a variety of functions, including detoxification, DNA repair, immunity, and oxidative stress. With some exceptions, these proteins are generally expressed more strongly in drones relative to workers, which, coupled with very low levels of hex110 (a major amino acid storage protein) in drones, suggests that drones favour broad, non-specific upregulation of putative stress response proteins to the detriment of retaining amino acid reserves. We speculate that this may improve their likelihood of protection against mild stressors but impair their ability to withstand prolonged or intense stress, since they have few reserved resources to draw on. Future research should focus on effects of multi-stressor exposures on drones and genetic variability of stress tolerance in the population, which are major gaps in knowledge of honey bee reproduction.

## Methods

### Drone and worker survival

In May, newly emerged (callow) drones and workers from three different colonies located in Vancouver, Canada were marked with paint pens and allowed to age for five days in their respective colonies. On Day 5, they were collected and placed in wooden California queen cages containing fondant. Each painted bee (both drones and workers) was placed in a separate cage with five other young (non-flying) worker bees, which were not analyzed in the experiment, to attend the drones. Different stress tests were conducted on different days, and the number of aged bees available differed for each sampling, but in all cases, there was roughly equivalent representation from each colony.

To analyze pesticide stress, bees were briefly anesthetized with carbon dioxide to immobilize them, then 2 µl of pesticide solution (either a cocktail mixture or imidacloprid, which was not part of the cocktail, in acetone) was applied directly to the thorax. The pesticide cocktail mixture (which contained tau-fluvalinate, coumaphos, 2,4-DMPF (2,4-Dimethylphenyl-N⍰-methyl-formamidine, a degradation product of amitraz), chlorothalonil, chlorpyriphos, fenpropathrin, atrazine, pendimethalin, and azoxystrobin) was produced exactly as previously described^26^ based on wax residue data published by Traynor *et al* ^*21*^. We tested doses at 0x, 0.33x, 2x, and 10x, where x is the median concentration in wax found in commercial colonies in the U.S. (see **Table 1** for components and their concentrations). The imidacloprid solutions were produced by serial dilution of the technical chemical acquired from Chem Service Inc. (West Chester, PA). We tested doses of 0, 1, 10, and 100 ppm. For all treatments, bees were allowed to recover for two days at room temperature in the dark, and were provided with two drops of water (∼100 µl) per day. In addition to the pesticide challenge, we also tested cold stress susceptibility. For the cold stress treatment, we placed the caged bees in a covered container in a 4 °C refrigerator for 0, 2, or 4 h and allowed all bees to recover as described above. After the two day stress recovery period, we counted the number of bees that were alive and dead. Workers and drones from the highest sublethal doses tested (10x cocktail, and 10 ppm imidacloprid) were euthanized by submerging in ethanol and then frozen at -20 °C for one week, until protein extraction. Cold stressed drones were not analyzed because survival was prohibitively poor.

**Table 1.**
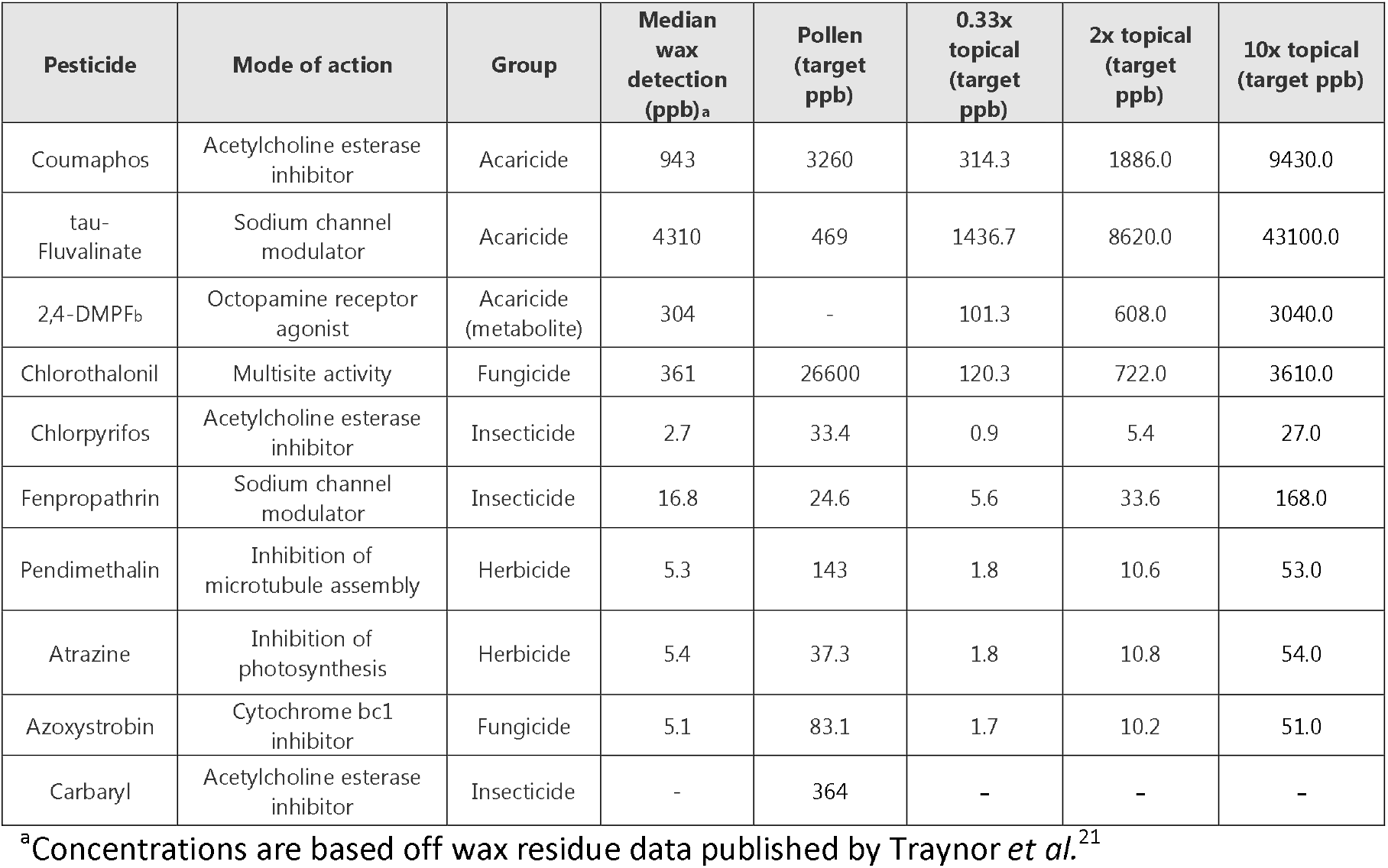
Target concentrations of the pesticide cocktail used for topical applications and pollen feeding

### Proteomics sample preparation

Hemolymph for the pesticide stress experiment was extracted from frozen bees by allowing them to thaw on ice, then using a scalpel to make a small incision between their 2^nd^ and 3^rd^ abdominal tergites. We inserted a small glass capillary against the incision to collect the hemolymph, then expelled the hemolymph (about 1-2 µl for workers, about 5-8 µl for drones) into 50 µl of 100 mM Tris buffer (pH 8.0). This solution was mixed and then spun at 16,000 x *g* to remove cellular debris, then transferred to a new tube. Clarified solution was precipitated using acetone (added 4 volumes of ice-cold acetone, incubated at -20 °C overnight). Precipitated protein was pelleted by spinning at 10,000 x *g* for 15 min, the supernatant was discarded, and the pellet washed with 500 µl of 80% ice cold acetone. The supernatant was discarded and the pellet allowed to dry at room temperature prior to suspending in digestion buffer (6 M urea, 2 M thiourea, 100 mM Tris, pH 8). Protein concentration was determined with a Bradford assay, and samples were prepared for mass spectrometry exactly as previously described^26^. Briefly, the samples were reduced, alkylated, digested with Lys-C, diluted with five volumes of 50 mM ammonium bicarbonate, then digested with trypsin overnight. Peptides were desalted using C18 STAGE tips^42^ as previously described, quantified via nanodrop, and analyzed on an Easy nLC-1000 (Thermo) connected to a Bruker Impact II Q-TOF mass spectrometer in randomized order.

Mass spectrometry data was searched using MaxQuant (v 1.6.1.0) with match between runs and label-free quantification enabled. The proteinGroups.txt file was imported to Perseus (v 1.6.1.1), where proteins only identified by site, potential contaminants, and reverse hits were removed, followed by proteins identified in fewer than three replicates of each group. This filtered the total proteins from 1,452 down to 654, which is the set upon which we performed our statistical analysis.

For the analysis of untreated drones and workers, newly emerged bees were marked with paint and allowed to age in their respective hives for 5 d. They were then retrieved and hemolymph was immediately extracted using glass capillaries. The hemolymph was then prepared for proteomics exactly as described above, except 200 ng of peptides were analyzed on a Bruker TIMS-TOF mass spectrometer and MaxQuant version 1.6.8.0 was used to process the data. Altered MaxQuant search parameters include: The precursor mass error for the main search was set to 50 ppm, and TIMS-DDA was selected within the “Type” tab of Group Specific Parameters, and LFQ and match between runs were enabled. The identified proteins were then filtered exactly as described above.

### In-hive pesticide treatments

Colonies were exposed to a pesticide cocktail via pollen patty feeding. The pesticide treatment mixture was based off of bee bread residue data from Traynor *et al*^21^. This mixture contained similar components as the topical application described above, with the exception that carbaryl was included and 2,4-DMPF was excluded, but with different relative proportions, owing to differences in detections in beebread versus wax. The cocktail added to beebread is described in detail in **Table 1**. Treated pollen patties received the pesticide mixture, while control pollen patties did not receive any added chemical treatment other than an equal amount of solvent. All pesticides (≥98% purity) were purchased from Sigma Aldrich Inc. or Chem Service Inc. Wildflower pollen (Glorybee Foods Inc. Eugene, Oregon) was powdered using a laboratory blender and was mixed with a sucrose solution to create a pollen supplement (final concentrations: 43% pollen, 43% granulated sucrose, 14% water by mass). Acetone, which quickly evaporates at room temperature, was used as the solvent for pesticide dilutions and comprised < 2% of the final diet. Components were thoroughly mixed in a stainless steel bowl using an electric hand mixer (Model 62633R, Hamilton Beach, Glenn Allen, VA) and 40 g portions were placed on individual wax paper sheets and sealed in plastic bags at -20 °C until use.

This experiment was carried out at the Lake Wheeler Honey Bee Research Facility (Raleigh, NC). Six colonies housed in single-deep standard Langstroth hive boxes roughly equilibrated for initial colony conditions (*e*.*g*., worker population, pollen and honey stores, and capped and uncapped brood) were fitted with pollen traps to encourage consumption of control and treated pollen patties. These colonies were continually exposed by placing a pollen patty on the top bars of each colony and consumption was recorded daily.

### Adult drone fostering experiment

This experiment was conducted in April-May with drones all sampled for analyses from May 3^rd^-7^th^. Frames of emerging drone brood were selected from two colony sources and allowed to emerge for 5 d in an incubator set to 33 °C and 55% RH. Emerged drones were collected daily in the morning and marked according to their source, placed into rearing cages and installed in one of six foster colonies treated with pollen patties as described above (three control, three treated). Foster colonies were fed either pesticide or control pollen patties for at least 28 d prior to the experiment. After 12 d had passed, drone cages were pulled from the colonies and the survivors were tallied and mean mass was taken. A subset of 20 drones from each cage were individually sampled for morphometric and reproductive measures.

### Drone larval rearing experiment

This experiment was conducted in May-June. Empty drone frames were placed into six experimental colonies (three treated and three control) once they had been fed control or treated pollen patties for at least 28 d. Only two of the pesticide-treated queens successfully reared drones. When the frames were capped and drone pupae had advanced to near-emergence (staged based on eye color), the frames were removed to an incubator set to 33 °C and 55% RH for emergence. Daily bees from each colony source were removed, marked and stocked into cages. All cages were fostered in a single, separate, untreated colony for 13 d post-emergence when they were removed from the colony and immediately returned to the laboratory for dissection and analysis. Drones from multiple emergence days and colony sources (but not treatments) were combined into single cages and paint-marked, with 7-73 drones contained per cage.

### Drone collection, dissection, and morphometric analysis

Drone collection and dissection proceeded according to previously established protocols^43^. Briefly, drones were weighed to the nearest 0.1 mg and anesthetized. Their head and thoraxes were photographed then dissected for their mucus glands and seminal vesicles removed and cut free from the testicles and ejaculatory duct; these were also photographed. Finally, the head, wings, and abdomen were cut free from the thorax and legs and these were weighed. For Experiment 1, the wings were not first removed from the thorax prior to weighing, therefore a corrective factor of 0.98 mg (the mean (N=5) of the four wings) was subtracted from their thorax masses for further analyses. The seminal vesicles were ruptured in 1.0 mL Buffer D and lightly homogenized^43-45^. Live and dead spermatozoa were visualized using the Invitrogen live/dead spermatozoa staining kit #L7011 (Carlsbad, CA) and read using a Nexcelom Cellometer® Vision Sperm Counter machine (Nexcelom Bioscience LLC; Lawrence, MA, USA) to gain a count of spermatozoa and viable proportion. The photographs were then analyzed using ImageJ version 1.51m9^46^ to measure the width of the head, thorax (as measured by the distance between tegulae), and the mean lengths of the seminal vesicles and mucus glands.

We defined body size as the first principal component of body mass, thorax mass, head width, and thorax width, and we defined fecundity as the first principal component of total sperm count, sperm viability (arcsine-transformed proportion), mean seminal vesicle length, and mean mucus gland length. These principal components represented 67.76% of the variation in their loading variables for body size and 40.87% of their loading variables for fecundity. We used these principal components to build linear models comparing the effects of colony source (a proxy for the combined effects of genetics and larval rearing environment) and adult rearing environment.

### Statistical analysis

Adult drone topical exposure and cold exposure survival counts were evaluated by logistic regression in R (v 3.6.0)^47^. Drones and workers were analyzed separately for each stressor using the generalized linear model glm(survival ∼ exposure, data = mydata, family = “binomial”), where survival is a binary variable (1 = survived, 0 = perished) and exposure is a continuous variable of either dose (for pesticide treatments) or duration of exposure (for cold treatments). Each stressor was analyzed separately. If a significant effect was identified, the model was rerun with exposure supplied as a categorical variable to identify the groups driving the significance.

All statistical analyses on the proteomics data were conducted in Perseus^48^ version 1.6.1.1 using simple two group comparisons and a Benjamini-Hochberg correction, which is better suited for high-throughput analyses of gene or protein expression than Bonferroni, at 5% FDR.

All analyses related to the hive exposures were performed in R (version 4.0.2), using ⍰^2^ tests for mortality. Linear models were conducted using linear mixed models, with parameter significances reported as a t distribution with estimated degrees of freedom or as a likelihood ratio test of the model with the parameter removed reported as a ⍰^2^. These analyses were performed using the lme4 package^49^. For experiment 1, where adults from two sources were installed into multiple treatment or control bank colonies, drone source colony and treatment were considered fixed effects and bank colony was considered a random, nested effect. For experiment 2, where larvae from control or treated colonies were reared to adulthood and banked in a common colony, treatment was considered a fixed effect and source a random effect.

## Data availability

Topical pesticide exposure and cold exposure survival data are available in *Supplementary Table S1*. Drone morphometric data are available in **Supplementary Table S2**. Survival data associated with colony exposures are available in **Supplementary Table S3**. Proteomics data of drones and workers exposed to acetone, imidacloprid, and cocktail treatments are available in **Supplementary Table S4**, and summary statistics of these data are available in **Supplementary Table S5**. Proteomics data of untreated workers and drones are available in **Supplementary Table S6**. All raw mass spectrometry data are available on MassIVE (www.massive.ucd.edu, accession: MSV000087818). Any code associated with this manuscript are available from the authors upon request.

## Author contributions

AM conducted the topical pesticide and cold exposure experiments, proteomics data acquisition, statistical analyses, and interpretation. JM and BM conducted the hive treatment experiments. BM conducted statistical analyses and interpretation of hive treatment experiments. JM produced the pesticide cocktails for topical and pollen exposures. Grants to LJF, DRT, and JM funded the research. AM wrote the first draft of the manuscript, with assistance from BM. All authors edited and approved the final version of the manuscript and contributed to the work intellectually.

## Acknowledgements

We would like to acknowledge Jason Rogalski and Renata Moravcova for running the mass spectrometry instruments, and Bradford Vinson and Jennifer Keller for helping to maintain the honey bee colonies.

## Funding sources

This work was supported by Genome Canada (264PRO), Genome BC, and BC Ministry of Agriculture grants to LJF. AM’s salary was supported by a fellowship from the Natural Sciences and Engineering Research Council. Funding support for BNM was provided by the USDA-NIFA project 2016-07962 and grant W911NF1920306 from the USARL. JPM was funded by a Graduate Student Fellowship through the NC Agricultural Foundation, a grant from the Foundation for Food and Agriculture Research (Grant #549053), and by the Foundation for the Preservation of Honey Bees.

## Notes

### Competing Interest Statement

The authors have declared no competing interest.

